# Multimodal MRI Insights into Glymphatic Clearance and Blood-Brain Barrier Integrity in Healthy Aging

**DOI:** 10.1101/2025.11.27.690842

**Authors:** Anjan Bhattarai, Yufei D. Zhu, Barah Albuhwailah, Pauline Maillard, Charles DeCarli, Audrey P. Fan

**Author notes:** Corresponding author: Anjan Bhattarai, Department of Neurology and Department of Biomedical Engineering, University of California Davis, 1590 Drew Avenue, Unit #100, Davis, CA 95618, USA.

## Abstract

Aging is associated with impaired CSF clearance in preclinical models, but its impact on blood-brain barrier (BBB) health and glymphatic function in humans remains unclear. We aimed to compare healthy younger and older adults using multimodal MRI to assess age-related changes in BBB permeability and glymphatic clearance, and to examine the relationships between these measures. Thirty participants were recruited (12 younger, 26 ± 3 years; 18 older, 75 ± 7 years). Diffusion-prepared arterial spin labeling measured the water exchange rate (Kw) across the BBB; free water (FW) imaging quantified extracellular free water in white matter; and diffusion tensor imaging along perivascular spaces (DTI-ALPS) assessed glymphatic function. Older individuals showed reduced whole-brain Kw (p=0.021), increased white matter FW (p=0.002), and reduced DTI-ALPS (p<0.001) compared to younger adults. Kw associated with both DTI-ALPS (β=0.402×10^−2^, p=0.022) and FW (β=-0.676×10^−3^, p=0.008). Reduced Kw may reflect impaired BBB function, while reduced DTI-ALPS and increased FW indicate impaired glymphatic clearance. FW partially mediated the relationship between Kw and DTI-ALPS, suggesting a potential mechanistic link. Overall, this study provides novel multimodal MRI insights into BBB and glymphatic alterations in healthy aging and may inform the development of MRI biomarkers to preserve cognitive health.

## Introduction

Healthy brain aging is a complex process which may involve subtle alterations in vascular, metabolic, and clearance pathways that maintain cerebral homeostasis^1^. One such pathway is the glymphatic system, which is defined as a brain-wide waste clearance system that facilitates the removal of interstitial solutes and metabolic waste, including β-amyloid (Aβ) and tau, from the brain tissue^2,3^. This system was first described in 2012 and was defined based on fluid transport pathway in mice^3^. It includes the flow of subarachnoid cerebrospinal fluid (CSF) into the brain’s interstitial spaces through para-arterial influx. Once inside the brain parenchyma, CSF mixes with interstitial fluid (ISF), which is then cleared through major veins (i.e., para-venous routes). This exchange is supported by the perivascular spaces and regulated by the neurovascular unit (NVU). The NVU includes components such as blood vessels, the blood-brain barrier (BBB) formed by endothelial cells and connected by tight junctions, perivascular spaces that are essential for CSF flow, and astrocytes that express aquaporin-4 (AQP4) at their endfeet^3,4^. These channels facilitate convective movement of CSF into interstitial spaces. The BBB protects the brain against ion fluctuations in the bloodstream and helps maintain optimal ISF conditions to preserve homeostasis^5^. The BBB and glymphatic functions are complementary and may partially overlap. However, their interaction is yet to be fully elucidated.

Impaired glymphatic clearance can be an initiating factor of age-related brain pathologies^2^. Preclinical studies indicate that glymphatic function declines with advancing age. In mouse models, impaired glymphatic function in the aging brain slows the clearance of interstitial Aβ, increasing the risk of developing neurodegenerative diseases^6^. Glymphatic dysfunction has also been associated with altered AQP4 expression, and changes in CSF influx, which depend on arterial pulsatility^6,7^. In parallel, increased BBB permeability has been reported in cognitively unimpaired older individuals^8^. Additionally, altered BBB permeability has been reported in the aging human hippocampus^9^. BBB transport and glymphatic clearance are interdependent^2^, and both are involved in removing interstitial solutes and maintaining fluid homeostasis. However, the mechanistic relationship between these functions in human aging has been poorly understood. Understanding this link could pave the way for developing MRI biomarkers to support cognitive health in aging populations.

Recent advances in MRI now allow us to assess BBB integrity and glymphatic function in vivo. Diffusion-prepared arterial spin labeling (DP-ASL), a non-contrast MRI technique, allows us to estimate the water exchange rate (Kw) across the BBB as a surrogate measure of BBB health^10^. Kw has been associated with a diffusion-weighted imaging (DWI)-based extracellular free water (FW) measure in older adults, indicating its role in CSF clearance^11^. Recent multimodal neuroimaging findings further indicate relationships between neurovascular integrity and fluid transport, including associations between FW and enlarged perivascular spaces, as well as between BBB Kw and cerebral blood flow (CBF)^12^. The mechanistic link between BBB Kw, FW, and glymphatic function remains unclear, providing the motivation for this study to investigate their contribution to glymphatic clearance in aging. The DWI-based technique, diffusion tensor imaging (DTI) along the perivascular pathway (DTI-ALPS) provides a surrogate measurement of glymphatic activity in the brain^13^. In this study, we aimed to compare BBB integrity, white matter free water, and glymphatic function between healthy young and older adults, and to assess the associations among these measures using multimodal MRI techniques, including DP-ASL, DTI-ALPS, and DTI-FW. These analyses may provide novel insights into glymphatic clearance and may guide interventions to preserve cognitive health in aging populations. We hypothesized that with aging, BBB function is reduced, leading to increased white matter free water accumulation and reduced glymphatic function.

## Methods

Ethics approval for this study was obtained from the UC Davis Institutional Review Board. All participants were at least 18 years old and provided written informed consent. Older participants were recruited through the UC Davis Alzheimer’s Disease Research Center (ADRC) longitudinal cohort and were at least 55 years of age. Older participants were evaluated to be cognitively normal within 12 months of the MRI scan through clinical evaluation by experienced neurologists at UC Davis Health and had no history of neurological disorder. Healthy, younger participants were recruited from an institutional registry of study volunteers.

### MRI data Acquisition

30 healthy participants with no known neurological or psychiatric disorders, including 12 younger (26 ± 3 years) and 18 older individuals (75 ± 7 years), underwent MRI scans on a 3T Siemens Prisma scanner. DWI, DP-ASL, and structural MRI data were acquired and processed as in **Figure 1**. Data were acquired with the following acquisition parameters: DWI with resolution = 2 x 2 x 2 mm^3^, 127 directions at b-values = 0, 500, 1000, and 2000 s/mm²; DP-ASL with post-label delay = 1800 ms, labeling duration = 1500 ms, resolution = 3.5 x 3.5 x 8 mm^3^, 3D GRASE readout, and diffusion-weighting with flow encoding arterial spin tagging (FEAST)^14^ at b-values = 14 s/mm² (to adjust for arterial transit times) and 50 s/mm². T1-weighted (T1W) magnetization-prepared rapid gradient-echo (MPRAGE) structural images were acquired with a resolution of 1.0 × 1.0 × 1.0 mm³.

**Figure 1:**
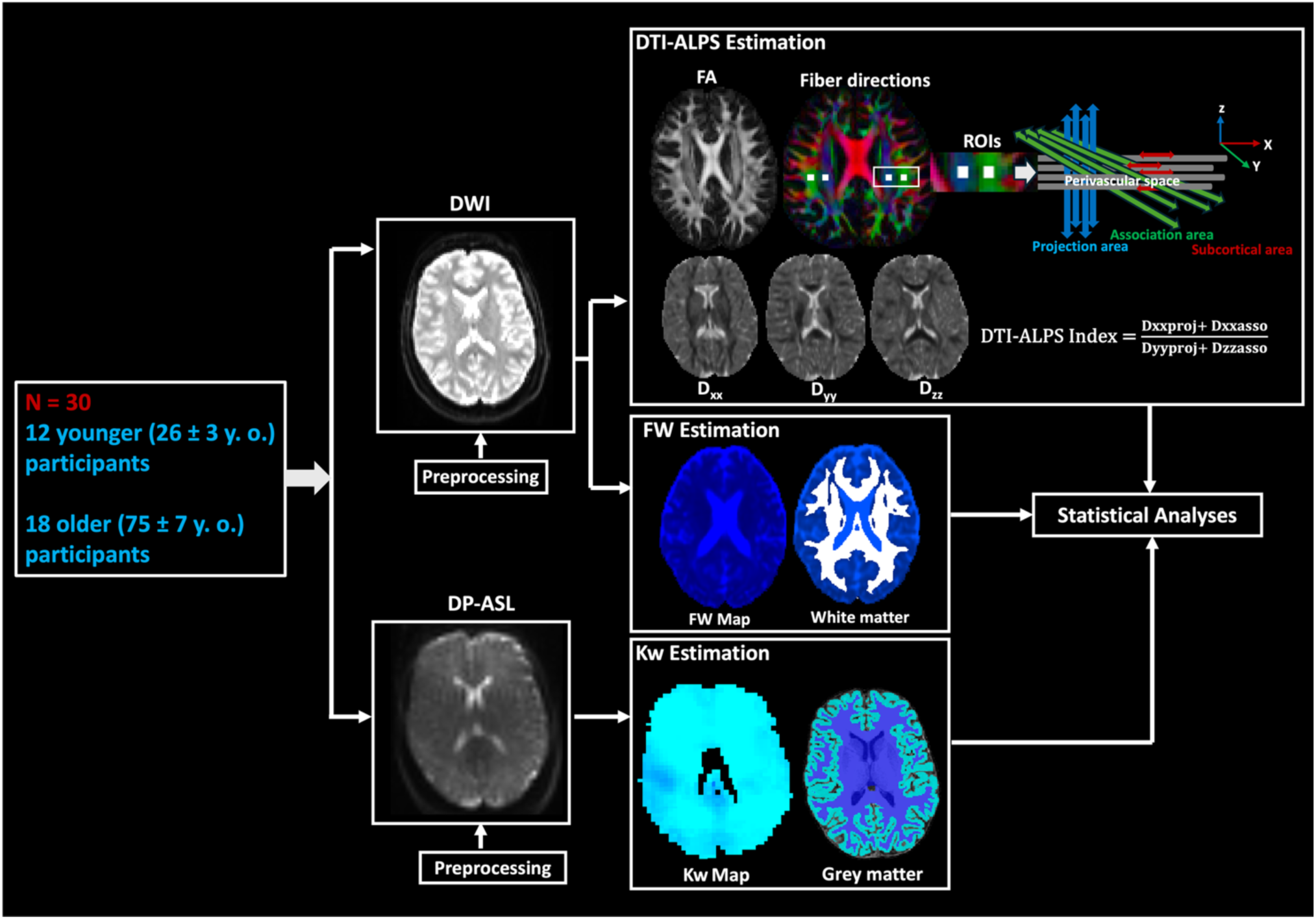
Summary of workflow and data processing. DWI: Diffusion-weighted imaging; DP-ASL: Diffusion-prepared arterial spin labeling; DTI-ALPS: Diffusion tensor imaging–analysis along the perivascular space; FA: Fractional anisotropy; Dxx, Dyy, Dzz: Diffusion coefficients; Kw: Water exchange rate.

### DP-ASL data processing

Raw DP-ASL data were processed to generate individual Kw images using the water exchange quantification (WEQ) toolbox with a two-compartment kinetic model^15^. The grey matter regions of interests (ROIs) were obtained using FreeSurfer cortical parcellation of the T1W images^16^. Kw was estimated for whole-brain, cortical grey matter, hippocampus, entorhinal cortex, and amygdala, with voxel-wise values averaged to obtain mean Kw (bilateral averaging applied to all regions except whole-brain).

### DWI data processing

DWI images were preprocessed using MRtrix3 and FSL software^17,18^. Preprocessing steps included denoising (dwidenoise), Gibbs ringing correction (mrdegibbs), motion and eddy-current correction with dwipreproc (FSL eddy, using topup for susceptibility-induced distortion correction), and bias field correction (dwibiascorrect)^18^.The preprocessed data were then used to fit diffusion tensors with FSL’s dtifit. ROIs (square, 3×3 voxels, 6×6 mm²)^19^ were manually placed in the projection and association areas along the perivascular space, guided by color-coded fractional anisotropy maps, and carefully inspected for anatomical accuracy. Bilateral individual diffusion coefficients along x y and z direction in the projection and association fibers and DTI-ALPS measures were estimated for each participant. Left and right DTI-ALPS measures were computed separately and then averaged to obtain a single ALPS index for each participant. The ALPS index was computed as:

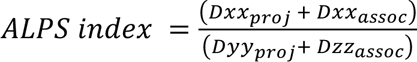; where Dxx, Dyy, and Dzz are the diffusion coefficients along the x, y, and z directions in the projection (proj) and association (asso) fiber ROIs, following the approach of Taoka et al^13^.

Preprocessed DWI data were modeled using a two-compartment bi-tensor approach, which separates the diffusion signal into tissue and isotropic free water contributions. This yielded voxelwise FW maps, reflecting extracellular water content ^20,21^. FSL’s FAST was used to segment T1-weighted images to obtain whole-brain white matter tracts^22^. Mean FW values were computed within the whole-brain white matter mask for each participant.

### Statistical analysis

Subjects were grouped by age (younger vs. older adults). Statistical comparisons between groups were performed using two-sample t-tests. Degrees of freedom, p-values, t-statistics, and 95% confidence intervals were calculated and reported. Group differences in Kw were assessed across the whole brain and in selected regions of interest (cortical grey matter, hippocampus, entorhinal cortex, and amygdala), with ROI-level analyses presented as exploratory.

Similarly, group differences in DTI-ALPS, individual diffusion coefficients, and white matter FW were computed. Pearson linear correlation was used to examine the relationship between DTI-ALPS and whole-brain white matter FW. Linear regression analyses were then performed to assess whether whole-brain Kw were associated with FW and DTI-ALPS.

A mediation analysis was conducted in MATLAB to test whether whole-brain Kw is associated with DTI-ALPS, mediated by FW (i.e., to assess whether FW explains part of this association). In an exploratory mediation analysis, lateral ventricular volume residualized for estimated total intracranial volume was included as a covariate to account for the potential effects of brain atrophy, following the approach described previously^11^. Statistical significance of the mediation effect was assessed using a bootstrapping approach with 5,000 simulations to compute 95% confidence intervals. All analyses were performed in MATLAB.

## Results

### DTI-ALPS

We observed reduced DTI-ALPS index in older individuals compared to younger individuals (p< 0.001), **Table 1**. Group differences were also evident in individual diffusion coefficients, with a significantly increased diffusion coefficient along the y-axis in the projection fibers (D_yy_proj) (p =0.011) in older participants. Although not statistically significant, reduced diffusion coefficient values were noted in older participants along the x-axis in the projection fibers (D_xx_proj) (p = 0.079), **Figure 2**.

**Figure 2:**
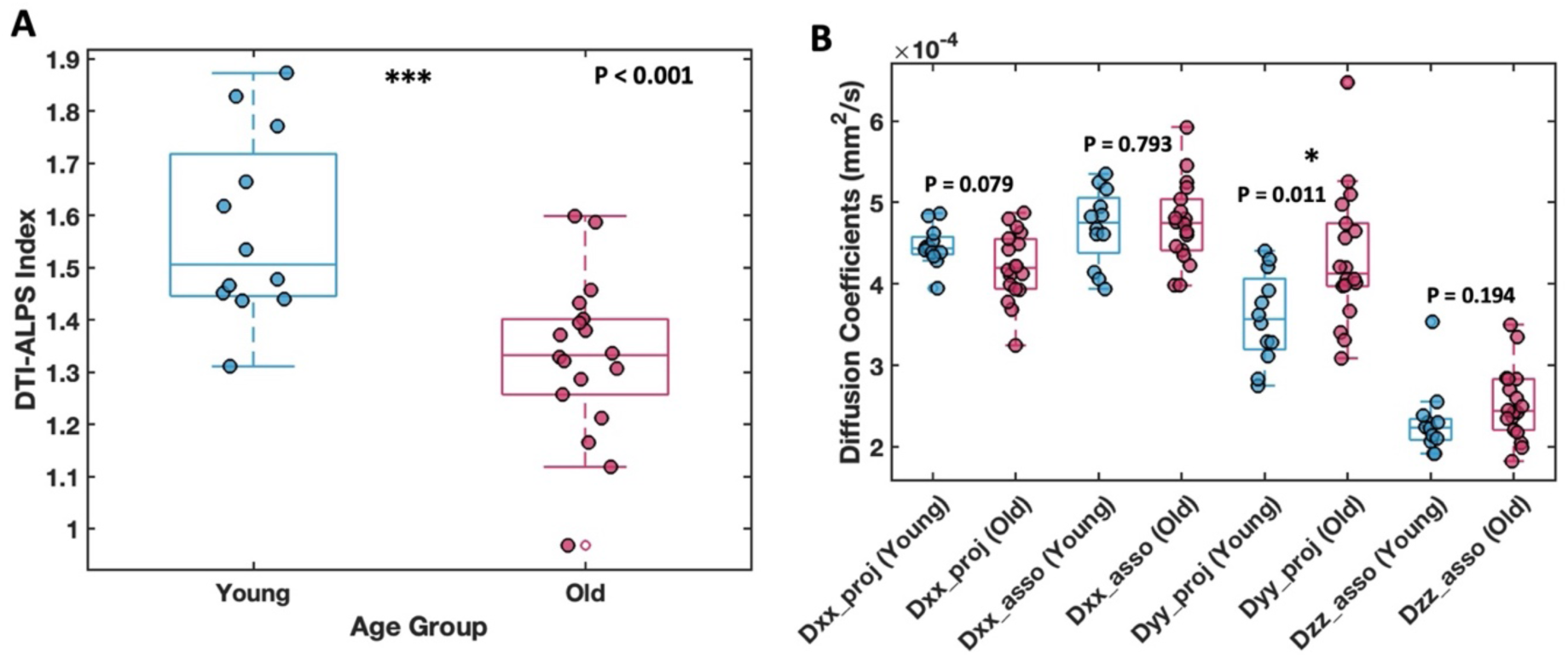
Group differences in glymphatic measures. A: DTI-ALPS index in younger and older participants. B: Group differences in individual diffusion coefficients.

**Table 1:**
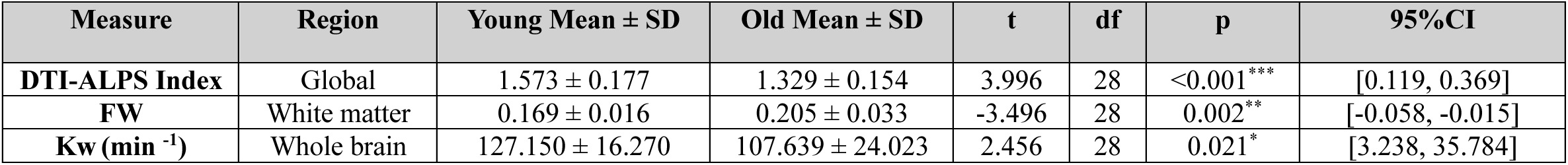
Whole brain/global multimodal MRI measures in younger and older adults.

| Measure | Region | Young Mean ± SD | Old Mean ± SD | t | df | p | 95%CI |
| --- | --- | --- | --- | --- | --- | --- | --- |
| **DTI-ALPS Index** | Global | 1.573 ± 0.177 | 1.329 ± 0.154 | 3.996 | 28 | <0.001*** | [0.119, 0.369] |
| **FW** | White matter | 0.169 ± 0.016 | 0.205 ± 0.033 | -3.496 | 28 | 0.002** | [-0.058, -0.015] |
| **Kw (min<sup>-1</sup>)** | Whole brain | 127.150 ± 16.270 | 107.639 ± 24.023 | 2.456 | 28 | 0.021* | [3.238, 35.784] |

### FW

Significantly increased white matter free water fraction was observed in older individuals compared to younger individuals (p=0.002), **Table 1**. Additionally, significant negative association (β = -3.487, p = 0.047) was observed between FW and DTI-ALPS measures across participants, **Figure 3**.

**Figure 3:**
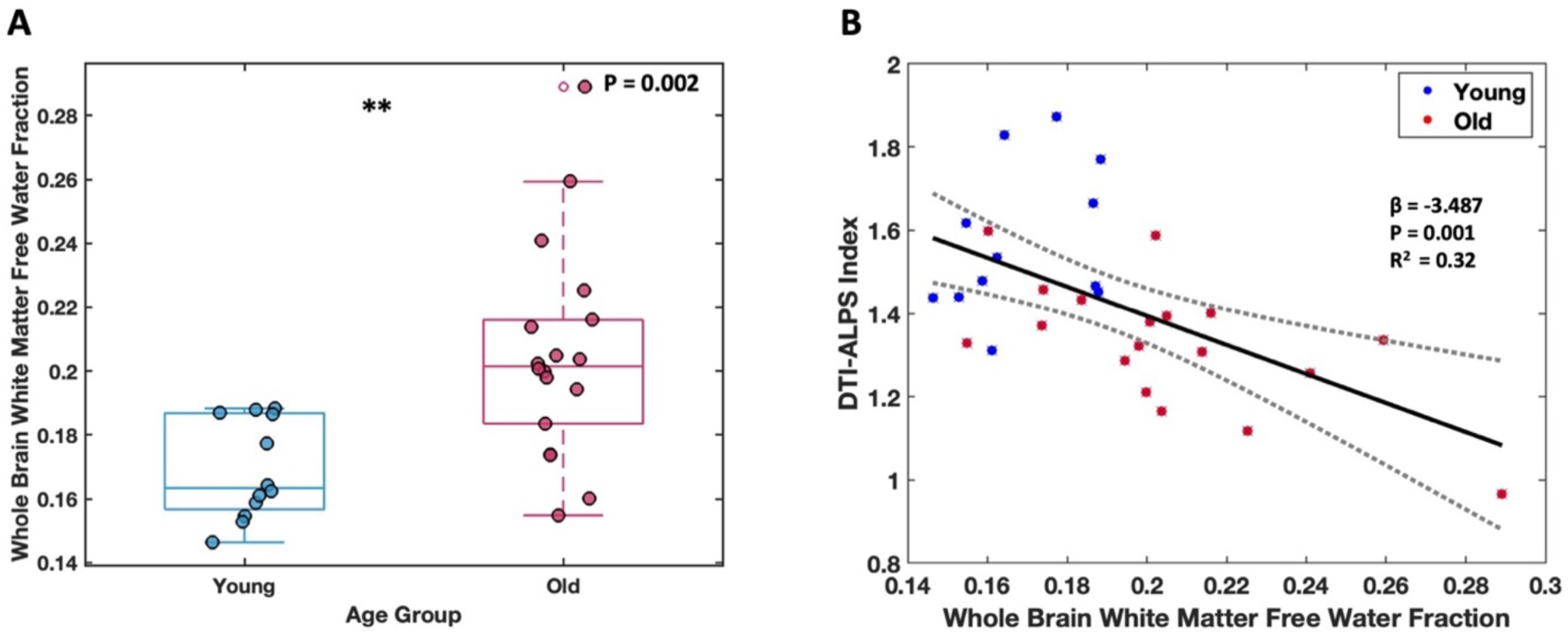
Group differences in FW measures. A: Whole-brain white matter FW fraction in younger and older participants. B: Association between whole-brain FW fraction and DTI-ALPS index.

### K_w_

Significantly reduced whole-brain K_w_ measures were observed in older participants (p = 0.021) **Table 1**, which negatively associated with mean FW (β = -0.676×10^−3^, p = 0.008) and positively associated with the DTI-ALPS index (β = 0.402×10^−2^, p = 0.022), **Figure 4**. Older participants had also significantly reduced K_w_ in cortical gray matter (p = 0.013). A reduction, though not reaching statistical significance, was observed in the hippocampus, entorhinal cortex, and amygdala, **Table 2**, **Figure 5**.

**Figure 4:**
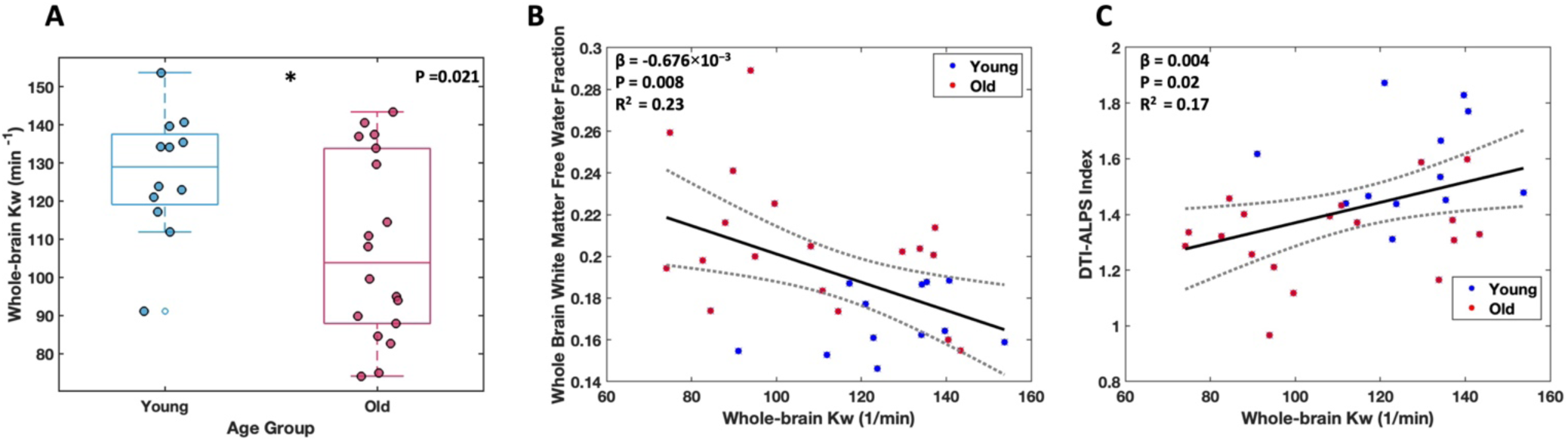
Group differences in whole-brain Kw (A) and its association with FW (B) and DTI-ALPS (C).

**Figure 5:**
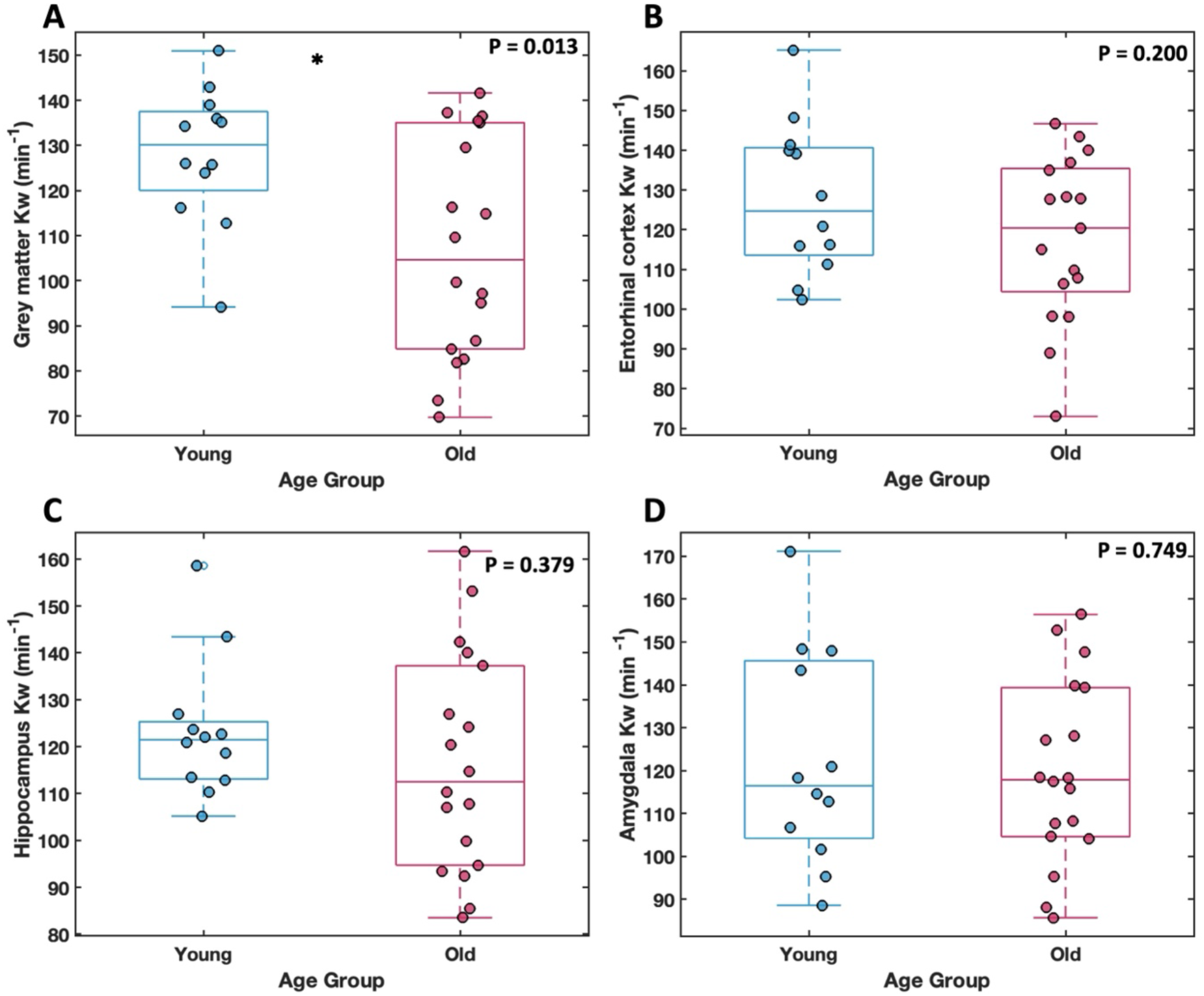
Group differences in Kw measures across grey matter regions: cortical grey matter (A), entorhinal cortex (B), hippocampus(C), and amygdala (D).

**Table 2:** Regional k_w_ measures in younger and older adults. P-values are uncorrected and presented to illustrate potential trends.

| Measure | Region | Young Mean ± SD | Old Mean ± SD | t | df | p | 95%CI |
| --- | --- | --- | --- | --- | --- | --- | --- |
| **Kw (min<sup>-1</sup>)** | Cortical grey matter | 128.078 ± 15.281 | 107.074 ± 24.480 | 2.641 | 28 | 0.013* | CI [4.710, 37.298] |
| | Hippocampus | 123.217 ± 14.771 | 116.400 ± 23.436 | 0.893 | 28 | 0.379 | [-8.813, 22.446] |
| | Entorhinal cortex | 127.816 ± 19.184 | 117.855 ± 20.728 | 1.313 | 27 | 0.200 | [-5.600, 25.520] |
| | Amygdala | 122.454 ± 24.989 | 119.709 ± 21.242 | 0.323 | 28 | 0.749 | [-14.651, 20.140] |

### Mediation analysis

We tested whether the effect of Kw on DTI-ALPS was mediated by FW. Mediation analysis revealed a significant indirect effect of Kw on DTI-ALPS index through FW (bootstrap mean *a*b = 0.0019, 95% CI [0.00014, 0.00404], p = 0.035; see **Figure 6**), indicating partial mediation.

**Figure 6:**
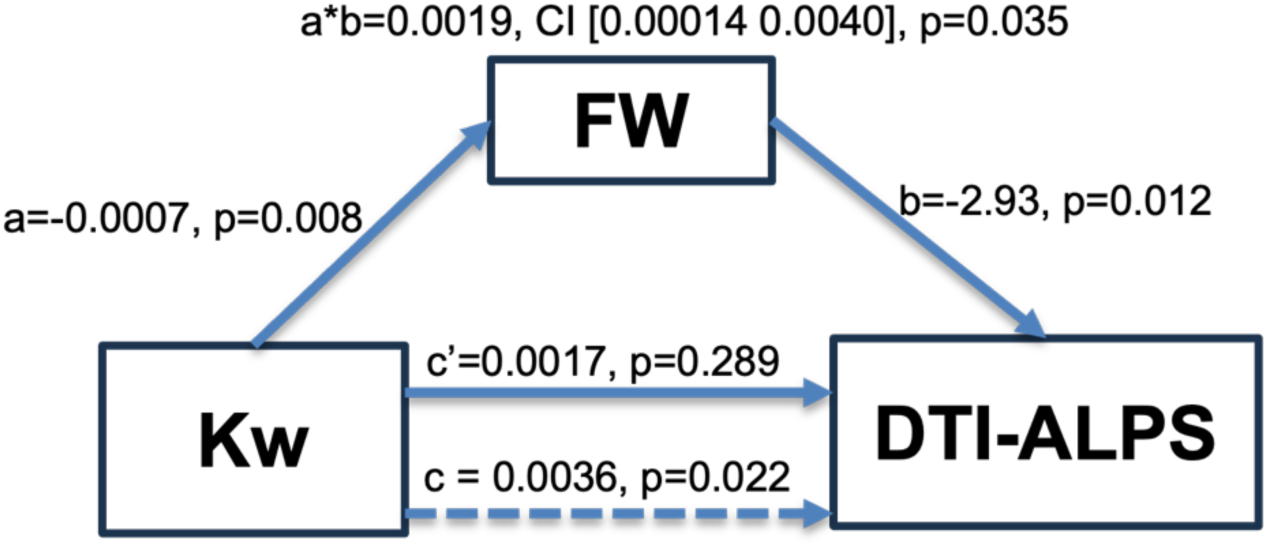
Mediation model showing the relationship between Kw and DTI-ALPS, partially mediated by FW. Arrows indicate the direction of effects, with coefficients labeled for both direct and indirect paths.

## Discussion

In this study, we used multimodal MRI to investigate the association between BBB integrity and glymphatic function in healthy aging. Our study cohort included healthy younger (26 ± 3 years) and older (75 ± 7 years) individuals, representing a clear age contrast to examine differences in BBB integrity and glymphatic function. We observed age-related changes in Kw, DTI-ALPS, and FW, suggesting declining BBB function and impaired glymphatic clearance in older individuals. In older individuals, Kw and DTI-ALPS were reduced, while white matter FW was elevated. Mediation analysis further indicated that FW partially explains the link between BBB integrity and glymphatic function, highlighting a potential mechanistic pathway connecting these processes.

Consistent with earlier findings, we observed a significant reduction in the DTI-ALPS index in older participants, suggesting a decline in glymphatic function with age^23,24^. This reduction accompanied by increased white matter free water, suggests that impaired perivascular CSF–ISF exchange may contribute to the accumulation of extracellular fluid. Reductions in whole-brain and regional Kw measures indicate subtle BBB dysfunction, which may further impede solute clearance. The trend of Kw reduction with aging is not yet fully understood. One potential explanation is dysregulation of water movement pathways due to age-related dysfunction of AQP4^25^. Subtle BBB dysfunction may permit ongoing water influx while reducing clearance efficiency, leading to increased FW. This hypothesis needs to be further explored. However, significant differences between younger and older Kw values reflect compromised BBB integrity in the aging process.

Our findings are consistent with earlier DP-ASL studies, which suggested a reduced Kw with aging^26^. However, Kw measured with multi-echo arterial spin labeling (ME-ASL) showed the opposite trend, increasing in older individuals. This difference in direction is not fully understood and may reflect differences in acquisition and modeling approaches between the two techniques. Dynamic contrast-enhanced MRI (DCE-MRI) is another commonly used technique to study BBB permeability^27^. An earlier study comparing K_trans_ (obtained with DCE-MRI) with Kw revealed that these measures were positively associated only in the white matter, caudate, and middle cerebral artery perforator territory, but not at the whole-brain level or in grey matter regions, suggesting that they may reflect different underlying BBB mechanisms^28^. Kw, which uses water as an endogenous tracer that crosses the BBB more readily than gadolinium contrast agents, also showed high test–retest reliability and reproducibility^28^.

Our results indicate that BBB integrity predicts impaired glymphatic clearance, potentially mediated by extracellular white matter FW. This suggests that BBB dysfunction may increase extracellular water content, which in turn reduces glymphatic clearance efficiency. This finding is conceptually consistent with an earlier study that also examined BBB integrity (i.e., Kw) and reported a mediating role of free water for cognitive outcomes in aging^11^. Previous findings in elderly individuals reported that BBB Kw is associated with CBF, along with links between FW, enlarged perivascular spaces, and white matter hyperintensities (WMH), highlighting interactions between NVU integrity and fluid regulation^12^. Our findings provide novel insights into the mechanistic role of NVU function (i.e., BBB integrity) in glymphatic clearance. Importantly, our findings highlight that subtle age-related shifts in BBB integrity and glymphatic function can occur even in the absence of disease.

While DTI-ALPS is a promising surrogate marker measure of CSF flow along the perivascular spaces, its sensitivity and specificity remain debated^29^. It can be compromised by the selection and placement of the ROI, as well as by the resolution of the DWI data and the processing approach. In our study, ROIs were manually placed to improve anatomical accuracy. DTI-ALPS has been increasingly used as a proxy for glymphatic function, and recent improvements such as DTI-ALPS plus may enhance its sensitivity^30^. Our study shows a clear distinction between younger and older DTI-ALPS measures, suggesting impaired glymphatic activity with age. Impaired glymphatic function may lead to the accumulation of metabolic byproducts over time, potentially compromising cognitive resilience ^31^. Sleep is known to enhance glymphatic clearance, and age-related changes in sleep may further influence CSF–ISF exchange efficiency^32–34^. These results need to be further validated using improved methods for assessing CSF clearance and standardized pipelines to ensure reproducible findings.

Conventionally, FW has been used to increase the sensitivity of traditional DTI metrics such as fractional anisotropy and mean diffusivity. More recently, it has gained attention as an individual surrogate marker of cerebral injury^35^. Our findings revealed significantly higher extracellular free water content in the white matter of older individuals, consistent with earlier findings, which may reflect impaired WM structural integrity in aging^36^. However, there is no direct evidence so far that excess FW content is related to neurodegeneration, which warrants further investigation. As hypothesized, we observed a significant correlation between Kw and both DTI-ALPS and FW measures, suggesting that cerebral fluid clearance mechanisms are interconnected. It is noted that in this study we focused only on WM free water content and not on gray matter structures.

Our mediation analysis suggests that impaired BBB integrity can predict glymphatic clearance through white matter FW accumulation. However, these findings were not significant when we included ventricular size as a covariate in the model. This suggests that the previously observed mediation of extracellular free water may, at least in part, reflect brain atrophy, as reflected by residualized ventricular volume. These findings may also be influenced by our relatively small sample size and should be validated in a larger cohort. Notably, BBB Kw measures were individually significantly correlated with both FW and DTI-ALPS measures, suggesting that the BBB plays a significant role in CSF clearance in the brain. Overall, our findings revealed an association between an NVU component and fluid clearance.

This study has some limitations. We used a multimodal approach to investigate BBB integrity and glymphatic clearance in healthy aging, with a cohort that had a clear age differentiation (younger: 26 ± 3 years, older: 75 ± 7 years). However, our sample size was relatively small. There were also some technical limitations. DP-ASL, used for Kw estimation, unlike DWI and T1W images, has poor spatial resolution, which may introduce partial volume effects in Kw measurements within smaller brain structures^37^. In this study, we examined whole-brain Kw in addition to regional analyses. As previously discussed, DTI-ALPS is a proxy measurement of glymphatic function and can be influenced by ROI selection and placement, as well as by the resolution of the DWI data. Future work should focus on improving the sensitivity and standardization of Kw, FW, and DTI-ALPS measures. Furthermore, we recommend that future studies validate our findings in a larger cohort.

## Conclusion

In this study, we utilized multimodal MRI metrics to study BBB integrity and glymphatic clearance in healthy aging. Our findings suggest that BBB integrity predicts glymphatic activity and extracellular free water accumulation in healthy aging. Significant reductions in Kw may reflect impaired BBB function, while reduced DTI-ALPS and increased FW accumulation may indicate impaired glymphatic clearance in older individuals. FW may partially explain the link between Kw and DTI-ALPS, highlighting a potential mechanistic pathway connecting these processes. The findings hold potential for advancing imaging biomarkers to preserve cognitive health in aging populations and should be validated in larger cohorts.

## Acknowledgements

We would like to thank all participants, their families, and the clinicians involved in this study. Without their help, this study would not have been possible.

## Author contributions

**Anjan Bhattarai:** Conceptualization, Data Curation, Formal Analysis, Investigation, Methodology, Software, Validation, Visualization, Writing - Original Draft, Writing -Review and Editing. **Yufei D. Zhu:** Data Curation, Investigation, Methodology, Software, Validation, Writing -review and editing. **Barah Albuhwailah:** Investigation, Writing -review and editing. **Pauline Maillard:** Software, Writing -Review and Editing. **Charles DeCarli:** Funding Acquisition, Project administration, Resources, Writing -review and editing. **Audrey P. Fan:** Conceptualization, Data Curation, Funding Acquisition, Investigation, Methodology, Project administration, Resources, Supervision, Writing -review and editing

## Statements and declarations

### Ethical considerations

Ethics approval for this study was obtained from the UC Davis Institutional Review Board.

### Consent to participate

All participants were at least 18 years old and provided written informed consent.

### Consent for publication

Consent to publish has been received from all participants.

## Data availability statement

Data will be shared upon reasonable request for research only after ethical approval for the specific project.

